# Determining the role of inflammation in the selection of JAK2 mutant cells in myeloproliferative neoplasms

**DOI:** 10.1101/079020

**Authors:** Jie Zhang, Angela Fleischman, Dominik Wodarz, Natalia L. Komarova

**Affiliations:** Department of Mathematics, University of California Irvine, Irvine, CA 92697, USA; Division of Hematology/Oncology, University of California Irvine, Irvine, CA 92697, USA; Department of Ecology and Evolutionary Biology, University of California Irvine, Irvine, CA 92697, USA

## Abstract

Myeloproliferative neoplasm (MPN) is a hematologic malignancy characterized by the clonal outgrowth of hematopoietic cells with a somatically acquired mutation most commonly in JAK2 (JAK2^*V* 617*F*^). This mutation endows upon myeloid progenitors cytokine independent growth and consequently leads to excessive production of myeloid lineage cells. It has been previously suggested that inflammation may play a role in the clonal evolution of JAK2^*V* 617*F*^ mutants. In particular, it is possible that one or more cellular kinetic parameters of hematopoietic stem cells (HSCs) are affected by inflammation, such as division or death rates of cells, and the probability of HSC differentiation. This suggests a mechanism that can steer the outcome of the cellular competition in favor of the mutants, initiating the disease. In this paper we create a number of mathematical evolutionary models, from very abstract to more concrete, that describe cellular competition in the context of inflammation. It is possible to build a model axiomatically, where only very general assumptions are imposed on the modeling components and no arbitrary (and generally unknown) functional forms are used, and still generate a set of testable predictions. In particular, we show that, if HSC death is negligible, the evolutionary advantage of mutant cells can only be conferred by an increase in differentiation probability of HSCs in the presence of inflammation, and if death plays a significant role in the dynamics, an additional mechanism may be an increase of HSC’s division-to-death ratio in the presence of inflammation. Further, we show that in the presence of inflammation, the wild type cell population is predicted to shrink under inflammation (even in the absence of mutants). Finally, it turns out that if only the differentiation probability is affected by the inflammation, then the resulting steady state population of wild type cells will contain a relatively smaller percentage of HSCs under inflammation. If the division-to-death rate is also affected, then the percentage of HSCs under inflammation can either decrease or increase, depending on other parameters.

## 1 Introduction

Myeloproliferative neoplasms (MPNs) are a group of hematologic malignancies characterized by clonal outgrowth of hematopoietic stem cells (HCSs) with somatically acquired mutations most commonly in JAK2 (JAK2V617F) (Campbell and Green, 2006; Baxter et al, 2005; James et al, 2005; Levine et al, 2005; Kralovics et al, 2005). These mutations result in cytokine independent growth of hematopoietic progenitors and consequently lead to an overproduction of myeloid lineage cells. Patients with early stage MPN can spontaneously progress to more aggressive neoplasms, such as myelofibrosis or acute myeloid leukemia (AML). The current therapeutic goals in MPN are to reduce the risk of blood clots and manage symptoms; no therapy other than bone marrow transplantation alters the natural history of MPN. Moreover, clinical trials in MPN focus on end stage disease, with minimal attention to patients with early stage disease. It is possible, however, that intervening in early stage MPN can make a meaningful impact on patient outcomes. An understanding of the conditions that promote the emergence of MPN as well as progression from early stage to late stage disease is vital for designing chemoprevention measures that could halt progression of disease in patients with early stage MPN or could prevent disease in healthy individuals at risk of developing MPN.

Recent research strongly indicates that inflammatory processes create an environment that promotes the selection of JAK2^617*F*^ mutant cells and that interference with these inflammatory processes can prevent the expansion of the mutant clones. Details of how this selection occurs, however, are not well understood.

Chronic inflammation has been linked to many different cancers, promoting cancer via multiple proposed mechanisms including induction of DNA damage and production of inflammatory cytokines that support growth of malignant cells, see e.f. Mantovani et al (2008). In certain cancers there is an established strong connection between inflammation and cancer, for example Helicobacter pylori infections in gastric mucosa-associated lymphoid tissue lymphoma, Hepatitis B or C infections in hepatocellular carcinoma (HCC) and inflammatory bowel disease in colorectal cancer (CRC) (Algra and Rothwell, 2012; Rothwell et al, 2012a,b). The anti-inflammatory agent aspirin has been found to prevent the development of colorectal, esophageal, gastric, biliary and breast cancer. Many mechanisms have been proposed for how inflammation promotes cancer, including induction of DNA damage and recruitment of inflammatory cells that support the growth of cancer cells. Chronic inflammation may also create an environment that is selectively advantageous for mutant neoplastic cells while negatively impacting normal counterparts.

In hematologic malignancies the differential effect of inflammation on normal versus neoplastic hematopoietic is likely a more important mechanism of cancer development. There is constantly a selection process for which hematopoietic stem cells contribute to blood production. Moreover, it has been hypothesized that chronic inflammation negatively affects HSCs and leads to their premature aging and exhaustion. HSC clones which have mutated in such a way as to avoid these suppressive inflammatory cues would therefore have a selective advantage, and dominate hematopoiesis as the normal HSC pool becomes decreasingly fit from the negative effects of chronic inflammation.

Chronic inflammation is a characteristic feature of MPN (Verstovsek et al, 2010; Slezak et al, 2009; Tefferi et al, 2011; Tyner et al, 2010) and not only drives many of the debilitating constitutional symptoms associated with the disease but also creates an environment which is highly favorable for growth of the JAK2^617*F*^ neoplastic hematopoietic stem cells. Interference with these inflammatory processes can prevent the expansion of the mutant clones (Fleischman et al, 2011). The JAK2^617*F*^ mutant hematopoietic progenitors are resistant to inflammation, while JAK2WT cells are suppressed by inflammation, possibly through the induction of apoptosis, quiescence, or reduced self-renewal/increased differentiation of HSC. Moreover, JAK2^617*F*^ mutant cells themselves produce inflammatory cytokines (most notably tumor necrosis factor-alpha, TNF) and also induce bystander normal cells to produce inflammatory cytokines (Kleppe et al, 2015). If the mutants are rare, the effect of this would be minimal.

An understanding of how the JAK2^617*F*^ HSCs are selected for is vital for our ability to intervene at an early stage, or even to prevent the disease. In this paper we present a class of mathematical models, from very general to more specific, that aim to describe the competition dynamics of mutant and wild type cells in the context of MPN. The goal is to play out several possible scenarios and explore the consequences of theoretical assumptions. For example, one can imagine that inflammation increases the differentiation rate of wild type HSCs. One the other hand, it is also possible that it may decrease their division rate or increase their death rate. What are the consequences of such changes on evolution? Is there a way to distinguish different mechanisms by observing the population dynamics?

This paper contributes to the large and growing theoretical literature on hematopoietic stem cell dynamics and related diseases, see e.g. Glauche et al (2007); Marciniak-Czochra et al (2009); Foo et al (2009); Stiehl and Marciniak-Czochra (2012); Goyal et al (2015). There are several mathematical papers focusing on MPN and related disorders. In Attolini et al (2010); Haeno et al (2009); Podlaha et al (2012), MPN was considered specifically, and it was determined theoretically that JAK2^617*F*^ being an early event in cancer progression is consistent with the data. In Kent et al (2013), mathematical modeling techniques were utilized to study the mechanisms of JAK2^617*F*^ subclinical clonal expansion and the transition to overt MPN. Papers by Traulsen et al (2010); Dingli and Pacheco (2011) studied the fate of somatic mutations at different levels of hierarchy and concluded that only mutations occurring in HSCs (and not in cells downstream from it) have a chance of producing a longterm disease. These papers however did not explicitly consider the role of inflammation in the evolutionary dynamics of JAK2^617*F*^ mutants.

Inflammation has been extensively modeled mathematically, see the collection of articles published in a special issue of Mathematical Biosciences, and also reviewed in Vodovotz et al (2009) and in Foteinou et al (2009). These models study various contexts, including infection and wound healing, see Voit (2009). In the present study we consider the role of inflammation in the context of cancer dynamics, and study its effect on the selection of malignant mutations in MPN.

## 2 Stem cells, differentiated cells, and evolutionarily stable strategies

We begin by presenting a very general framework of thinking about hematopoietic stem cells (HSCs) and their niche, and trying to identify the role of inflammation in the rise of mutants.

### 2.1 Axiomatic modeling

Let us suppose that the population of interest consists of HSCs, *x*, and their more differentiated progeny, *y*. We assume that the dynamics of homeostatic tissue renewal in the HSC niche is controlled by factors related to the niche and also by factors secreted by the cells in the lineage (autonomous control). In particular, we will concentrate on the influence of inflammation on HSC dynamics. In a very general sense, tissue renewal is governed by the equations

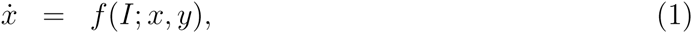

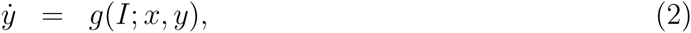

where *I* stands for the level of inflammation, which can directly impact relevant processes in the system. The following assumptions are readily made on the functions *f* and *g*:

1. *f* and *g* are continuous functions;
2. Stem cells are only produced by stem cells (no de-differentiation is assumed): *f* = 0 if *x* = 0;
3. *g* = 0 if *x* = *y* = 0 (no production of differentiated cells is assumed from cells other than *x* and *y*). Because of these requirements, it is convenient to rewrite *f* (*I*; *x, y*) = *γ*(*I*; *x, y*)*x* and *g*(*I*; *x, y*) = *α*(*I*; *x, y*)*x* + *β*(*I*; *x, y*)*y*, so we have

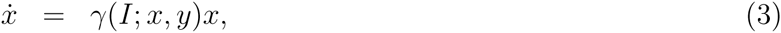

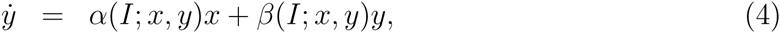 Functions *γ*, *α*, and *β* have units of inverse time. Here, *γ* describes the net growth rate of stem cells, which may comprise proliferation of stem cells, differentiation of stem cells, and their death. Rate *α* describes the production of differentiated cells by means of symmetric and/or asymmetric divisions of stem cells, and *β* describes possible divisions and deaths of the differentiated cells. The dependence of these three functions on the populations of cells indicates the presence of autonomous control. Assuming that differentiated cells cannot maintain themselves in the absence of stem cells, we set
4. *β*(*I*; *x, y*) < 0 (differentiated cells die in the absence of HSCs). It follows that *x* = *y* = 0 is a steady state. It is unstable if we postulate
5. *γ*(*I*; 0, 0) > 0 (stem cells can expand from low numbers). We will further assume that there exists another steady state, denoted as 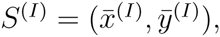, where 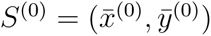 stands for the steady state in the absence of inflammation. We have

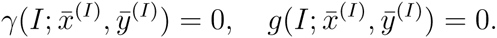 For stability of the nontrivial equilibrium, we require that

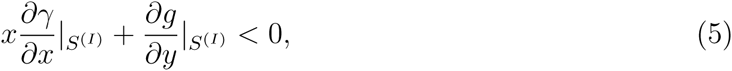

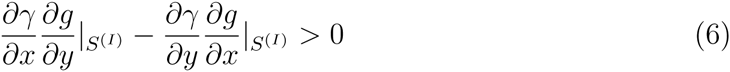

(these derive from the standard conditions that follow from the requirement that the real part of the eigenvalues of a 2 *×* 2 system is negative). The following assumption guarantees the former inequality:
6. Negative control: 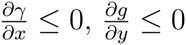 (but not simultaneously zero).

Conditions (**1-6**) above specify the reasonable mathematical assumptions that we can impose on the (generally unknown) biological rates. Narrower classes of functions and some specific examples are discussed later in the paper.

### 2.2 Mutants, growth, and equlibria

Next, we introduce a population of mutants that compete with the wild type population. Denoting by *x*_1_ and *y*_1_ mutant HSCs and differentiated cells, we have the following system describing the composite population:

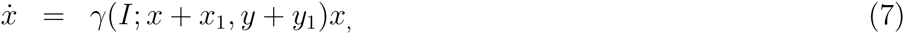

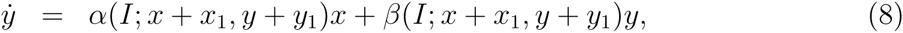

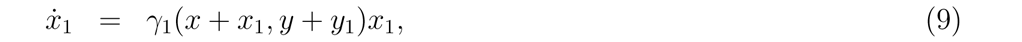

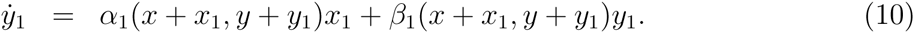

Here, *γ*_1_*, α*_1_, and *β*_1_ are the mutant equivalents of the corresponding wild type functions. Note that their dependencies of the population variables describe control and thus the total numbers of HSCs and differentiated cells (mutant and non-mutant) appear as their arguments. We do not include the process of de-novo mutation, and therefore populations *x*_1_ and *y*_1_ are not produced by populations *x* and *y*. We assume that while the wild type cells are influenced by the inflammation, the mutants are not. Therefore, functions *γ*_1_*, α*_1_, and *β*_1_ do not contain the argument *I*. We suppose that assumptions **1** *−* **6** above apply to mutant cells as well as the w.t cells. We further assume that in the absence of wild type cells, mutants are capable of stable homeostasis, that is, conditions similar to inequalities (5,6) apply to the mutant functions.

One important difference between mutants and w.t. cells is their growth rate in the absence of competition (that is, starting from low numbers, in the absence of the other population). These are given by

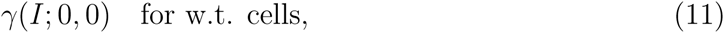

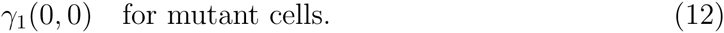

Comparing these two quantities will allow one to determine which population grows faster from low numbers, which is important for the analysis of repopulation experiments.

Next, we turn to the analysis of homeostatic conditions. In general, assuming phenotypic differences between w.t. and mutant cells, the following equilibria are possible in this system:

- The trivial equilibrium, *x* = *y* = *x*_1_ = *y*_1_ = 0; it is unstable due to condition **5**;
- The w.t. equilibrium, 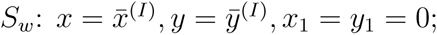
- The mutant equilibrium, 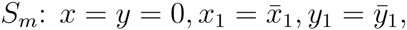 where

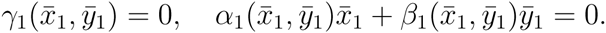

Next we investigate stability of equilibria *S_w_*and *S*_*m*_. For equilibrium *S_w_*, we notice that the Jacobian has the following structure,

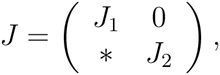

where *J*_1_ is the same as comes up in the stability of the w.t. cells in the absence of mutants and leads to conditions (5,6), and

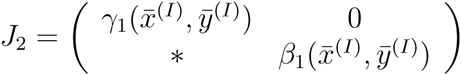

Because *β*_1_ < 0, we have the stability condition for the *S*_*w*_ equilibrium:

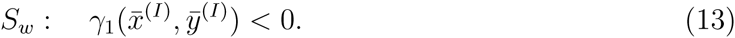

Similarly, the stability condition for the mutant equilibrium is

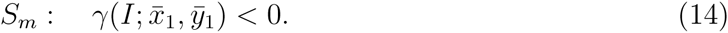

In other words, for equilibrium *S*_*m*_, stability is guaranteed if (i) mutants are capable of maintaining stable homeostasis in the absence of wild the cells, which is given by conditions similar to inequalities (5,6), and (ii) the mutant strategy is an ESS, that is, the competing strategy (the wild type) cannot invade from low numbers, as guaranteed by condition (14).

In this very general setting, we can make further progress by examining the effect of inflammation on the growth rate of HSCs. Given that the mutant equilibrium is stable in the presence of inflammation (condition (14)) and unstable in the absence of inflammation, which is equivalent to condition

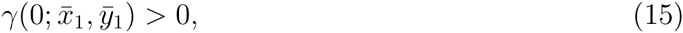

we can see that inflammation decreases the net growth rate *γ* of stem cells (at least, at point 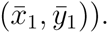. It is therefore reasonable to assume that inflammation decreases the net growth rate in a region of the (*x, y*) plane.

These concepts are illustrated in figure (1). In the figure, the functions *γ*(*I*; *x, y*), *γ*(0; *x, y*) and *γ*_1_(*x, y*) are presented, which are the growth rate of w.t. HSCs in the presence and in the absence of inflammations, and the growth rate of mutant cells, respectively. The functions are shown as a contour plot, where the axes are *x* and *y* and the three lines identify the sets of points where these functions are zero. We assume that the functions are positive below and negative above these lines. Let us first assume that no inflammation is present in the system. Then the point 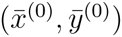 is a stable equilibrium, because the function *γ*_1_ is negative at this point (condition (14) with *I* = 0). This follows from the fact that point 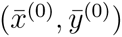 is above the *γ*_1_ = 0 line. Next, assume that inflammation is induced in the system. Then the growth rate of the w.t. cells is described by the *γ*(*I*; *x, y*) = 0 line and the corresponding equilibrium is 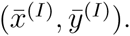. It is however unstable and instead, the mutant equilibrium 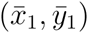 gains stability, because condition (14) is satisfied (the point 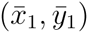 is above the *γ*(*I*; *x, y*) = 0 line).

**Figure 1:**
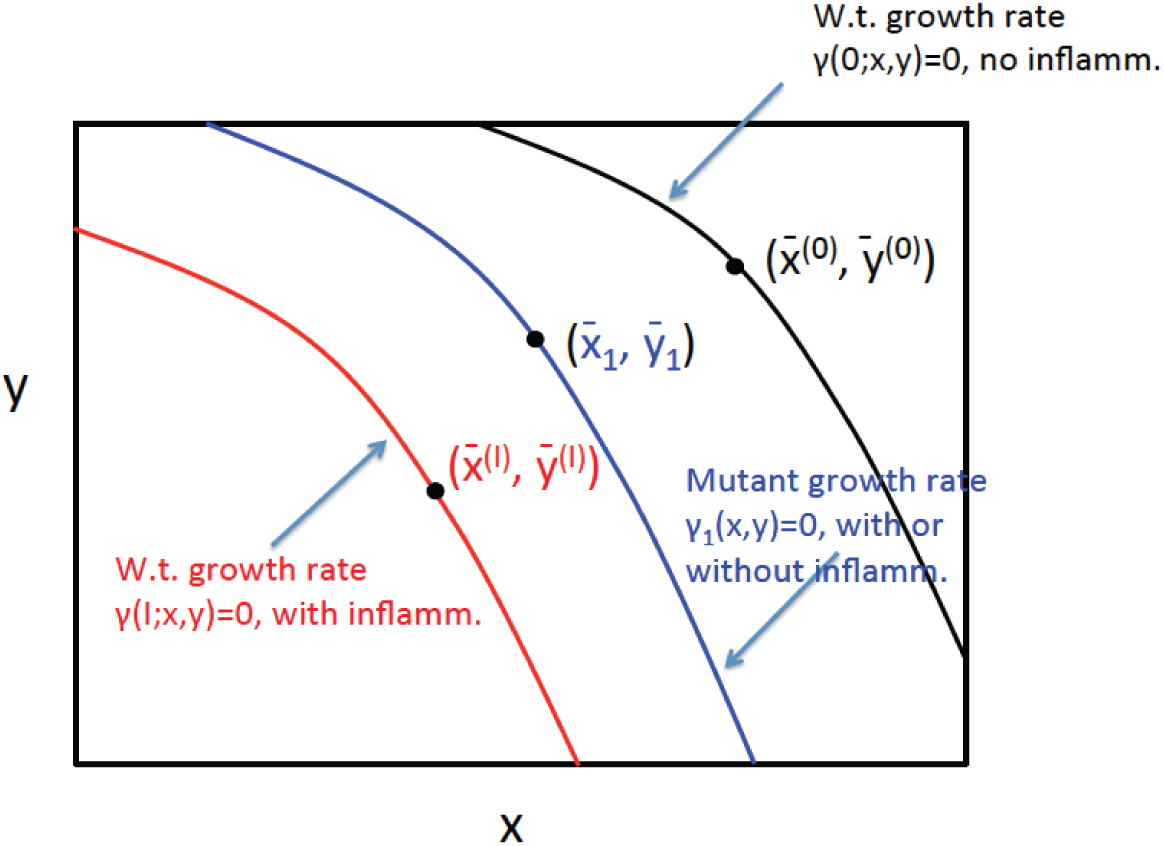
The concept of ESS in the context of w.t. and mutant cells in the presence and in the absence of inflammation. The functions *γ*(*I*; *x, y*), *γ*(0; *x, y*) and *γ*_1_(*x, y*) are presented by means of their contour plots, where the axes are *x* and *y* and the three lines identify the sets of points where these functions are zero. The equilibrium points are marked for each line. The positions of these points are determined by the intersection of the given lines with the lines *g*(*I*; *x, y*) = 0*, g*(0; *x, y*) = 0, and *g*_1_(*x, y*) = 0 respectively (not shown).

## 3 A stem cell control model with symmetric divisions

Next we consider a more concrete stem cell model, where the net growth rate of SCs is comprised of three processes:

- SC proliferation events, whereby SCs divide into two daughter cells each of which is a SC,
- SC differentiation events, whereby SCs divide into two daughter cells each of which is a differentiated cells,
- SC death.

The rates of these events can be expressed as follows. Denote by *R*(*I*; *x, y*) the total division rate of SCs and by *P* (*I*; *x, y*) the probability that upon division, the SC proliferates. Then the rate of proliferation events is given by *RP* and the rate of differentiation events is given by *R*(1 *− P*). The death rate of SCs is denoted by *D*(*I*; *x, y*), such that the net growth rate is given by

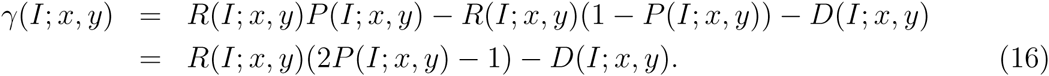

We assume that differentiated cells do not proliferate, but they can leave the compartment by death or further differentiation at rate *δ*(*I*; *x, y*). All the rates in general may depend on both (total) cell populations and on the level of inflammation. We have the system:

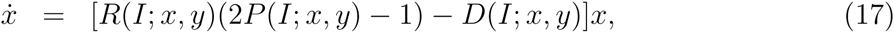

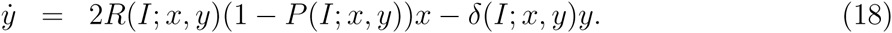

Apart from the trivial equilibrium, there is a nontrivial steady state, 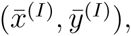, which is defined implicitly as the solution of the following two equations:

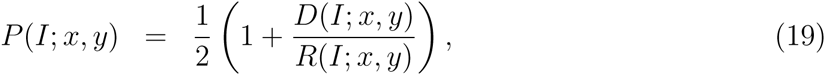

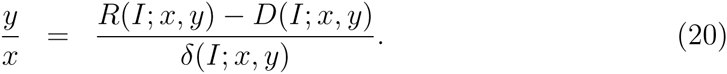

We will assume that this equilibrium, as well as the equilibrium corresponding to the mutants in the absence of w.t. cells, are stable, see conditions of type (5-6).

### 3.1 Mutant dynamics

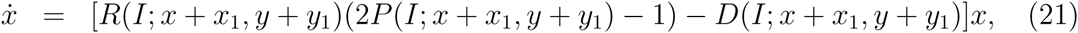

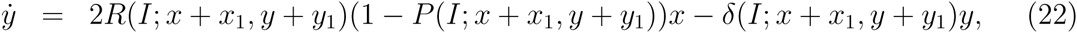

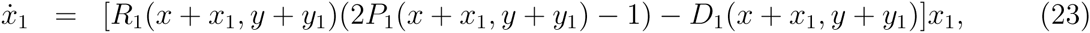

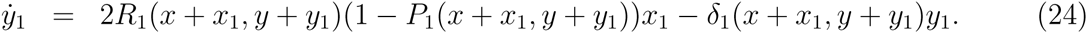

In this system, functions *R*(·) and *R*_1_(·), are growth rates of wild type and mutant HSCs respectively; functions *P* (·) and *P*_1_(·) are proliferation probabilities of wild type and mutant HSCs respectively; *D*(·) and *D*_1_(·) are death rates of wild type and mutant HSCs respectively, and *δ*(·) and *δ*_1_(·) are death (or exit) rates of wild type and mutant differentiation cells respectively. All these are functions the total number of HSCs and differentiated cells; in addition, the functions for the w.t. cells also depend on the inflammation level.

### 3.2 Growth rates in the repopulation experiments and the equilibria

The linear growth rates are again obtained by linearization of the equations, see also expressions (11-12):

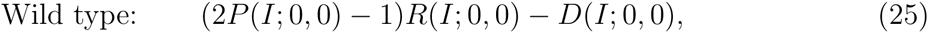

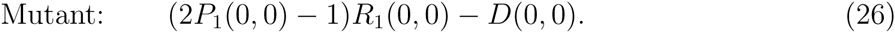

Next, we turn to homeostatic conditions. As before, we have three equilibria: the trivial equilibrium, and two nontrivial equilibria, which are defined implicitly as follows: the wild-type domination equilibrium,

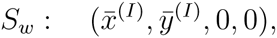

where the w.t. populations are defined by equations (19-20); and the mutant equilibrium,

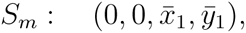

where the mutant populations are defined by equations similar to (19-20) except all the functions have subscript 1 for mutants.

Stability conditions follow from the ESS conditions (13) and (14) respectively. For example, to ensure that the mutant equilibrium is stable against the invasion of the w.t. cells in the presence of inflammation, we have to require that

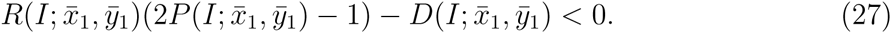

In the absence of inflammation, this equilibrium is unstable, such that

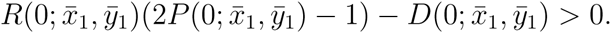

### 3.3 The effect of inflammation

As was established in the previous section, the effect of inflammation on the net growth rate of HSCs is to decrease it from a positive to a negative quantity. In the present model, we can say that this can be achieved through three mechanisms:

- Inflammation reduces the proliferation probability (thus increasing the differentiation probability) of HSCs;
- Inflammation decreases the division rate of HSCs;
- Inflammation increases the death rate of HSCs.

There are several interesting consequences of these arguments.

**In the absence of significant HSC death rates, the only effect of inflammation compatible with the rise of mutants is an increase in the probability of HSCs to differentiate.** If *D* = 0, the expression in (27) can only be made negative by decreasing the probability of HSC proliferation below 1/2,

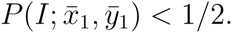

A decrease in the division rate *R* cannot ensure mutant stability in the presence of inflammation. At the same time, a decrease in *R* can lead to the decrease in the linear rate of HSC expansion, as following from expression (25).

**In the presence of HSC death, an alternative mechanism is a decrease in the divisions-to-death ratio,** *R/D*. Such a decrease can not only ensure mutant stability but also slow down the linear expansion rate of HSCs, expression (25).

The question is whether one can determine whether it is a decrease in the proliferation probability or a decrease in the divisions-to-death ratio that is responsible for the observed behavior. It is instructive to consider the equilibrium population values in the absence and in the presence of inflammation.

Suppose there are no mutants in the system. At equilibrium, we have equations (19-20), where the latter is the differentiated-to-HSC ratio. Let us assume that the rates are controlled by only one of the populations. For example, we can let *R*(*I*; *y*), *P* (*I*; *y*) be decreasing functions of the differentiated cell population, as was assumed in Rodriguez-Brenes et al (2011). We further suppose that *D* and *δ* are nondecreasing functions of *y*.

We first explore the scenario where the only effect of inflammation is the reduction in the probability of proliferation, *P* (such that *R*, *D*, or *δ* do not depend on *I*). For a fixed level of inflammation, the left hand side of equation (19) is a decreasing function of *y* while the right hand side is an increasing function of *y*. Increasing *I* from zero to a positive value, we shift the left hand side down, thus moving the intersection point to the left. Therefore, 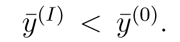. Next, from equation (20) we observe that the right hand side only depends on *I* through *y*^(*I*)^. A decrease in this equilibrium point will cause an overall increase in the right hand side, suggesting that the differentiated-to-HSC ratio increases with inflammation. Since both the value of *y* at the equilibrium and the *x/y* ratio decrease under inflammation, it follows that the total population size decreases.

Next, let us assume that the *R/D* ratio decreases with inflammation. As before, we turn to equation (19), whose right hand side now shifts upward, again resulting in a decrease in the value of *y* at intersection. As before, this in turn can increase the right hand side of equation (20). On the other hand, quantity *R − D* now directly decreases with *I*. Therefore, the situation is now more complicated, and it is not possible to say in which way the two opposing effects may trade-off. We conclude that direct dependence of the divisions-to-death ratio on inflammation will cause a decrease in the differentiated cell enrichment effect at equilibrium and can even reverse it. While the population of differentiated cells decreases under inflammation, we cannot determine the trend in the HSC population.

Similar analysis can be carried out if we assume that the rates are decreasing functions of the HSC population *x*. If the only effect of inflammation is to decrease the proliferation probability of HSCs, then it follows that the number of HSCs at equilibrium is smaller under inflammation, and the HSC-to-differentiated ratio is also smaller. We cannot determine whether the total population increases or decreases. In the case where the *R/D* ratio decreases with inflammation, we can only conclude that the total number of HSCs decreases under inflammation, but cannot argue about the HSC-to-differentiated ratio or the total population size.

### 3.4 Particular form of control

Let us consider a particular functional form of the controls in system (21-24):

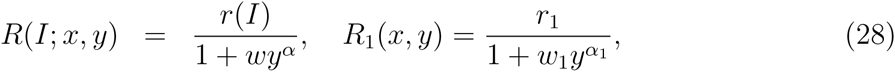

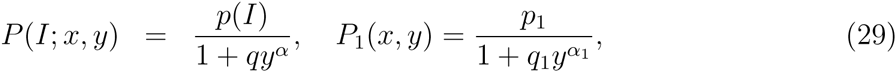

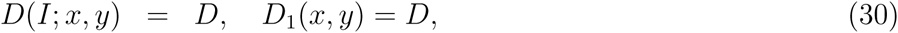

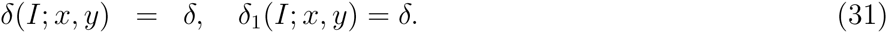

Here parameters *w*, *w*_1_, *q*, and *q*_1_ determine the strength of control; together with parameters *r*, *r*_1_, *p*, and *p*_1_ they define the magnitude of the division rate and the probability of proliferation. We are interested in two questions:

- Under what circumstances will mutant cells have a faster linear growth rate, when the population is expanding from low numbers?
- Under what circumstances will mutant cells come to dominate the population at homeostasis?

As we already know from the general system, the conditions for the two types of “domination” are quite different. We can model expanding populations by equations (21-24) where initially all the variables are very small compared to any nontrivial equilibrium values (see below for the quantification of those). The behavior at homeostasis can be studying by perturbing the system near an equilibrium.

We first consider expanding populations. Linearizing equations (21) and (23) with respect to small initial values of *x*, *y*, *x*_1_, and *y*_1_, we obtain the following expressions for the linear growth rates:

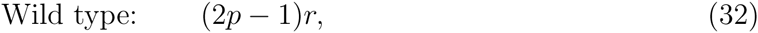

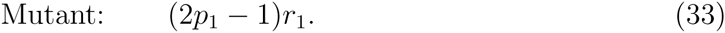

Next, we turn to homeostatic conditions. For analytical tractability let us assume that *α* = 1, but the results will also hold qualitatively for different values of *α*. We will further assume that the death rates of HSCs are negligible, and set

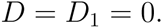

There are three equilibria that exist in system (21-24). The trivial equilibrium corresponds to population extinction:

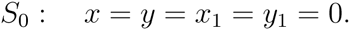

There is also an equilibrium corresponding to wild type excluding mutants:

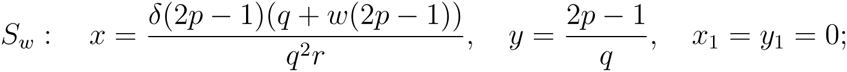

and the last equilibrium corresponds to the mutants excluding the wild type population:

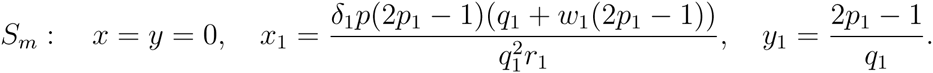

The first, trivial, equilibrium is positive and stable if *p*_1_ < 1/2 and *p* < 1/2. The second (mutant extinction) equilibrium is positive and stable if *p >* 1/2 and any of the following conditions hold: (ia) *p*_1_ < 1/2 or (iia) *p*_1_ > 1/2 and

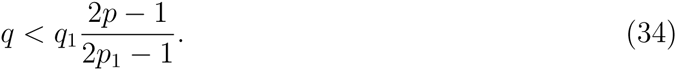

Finally, the third equilibrium is stable (that is, the mutants dominate) if *p*_1_ > 1/2 and any of the following conditions hold: 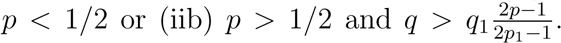 All the conditions are summarized in figure 3.

**Figure 3:**
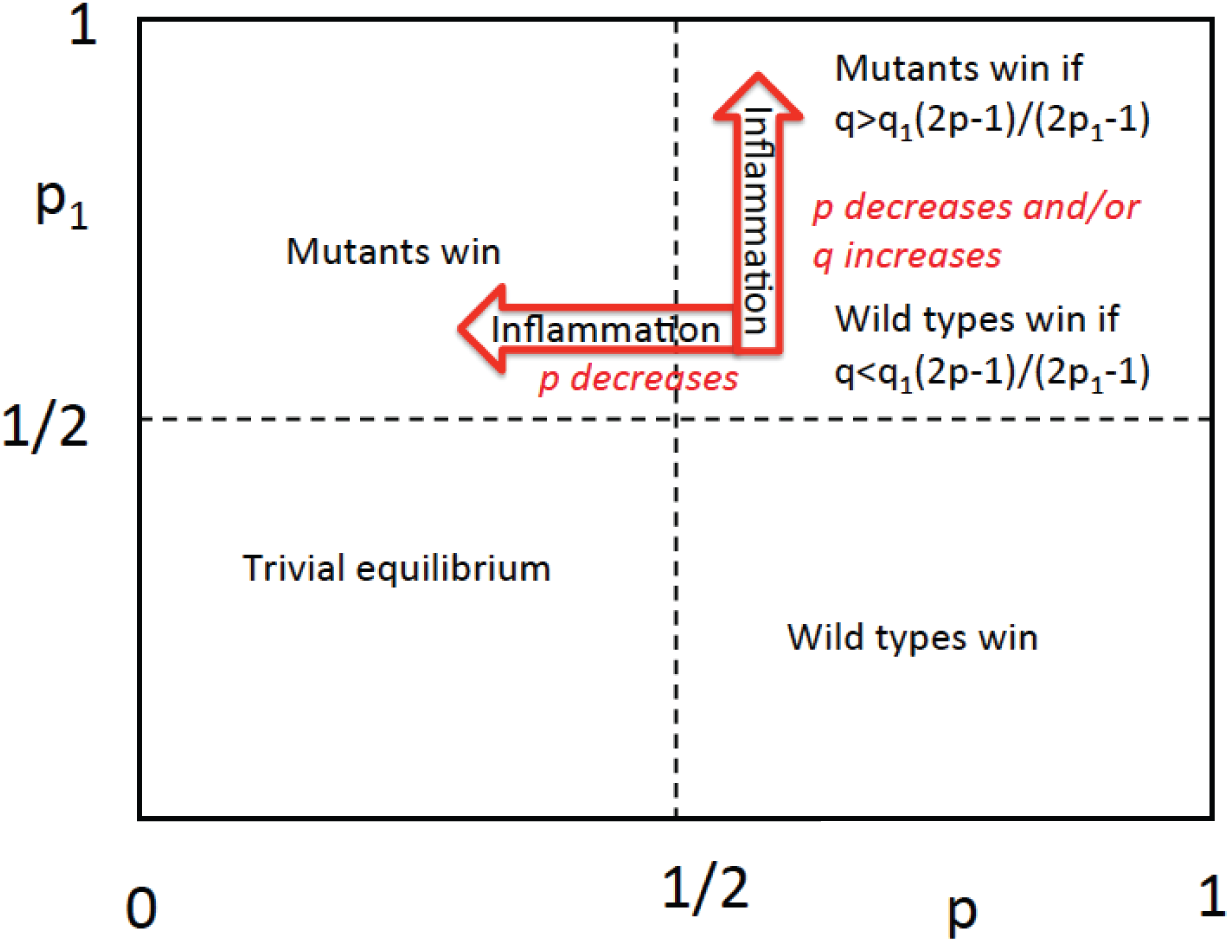
Theoretical prediction for the dominance in homeostasis in model (21-24) with functional forms given by (28-31). The horizontal axis is *p* and the vertical axis is *p*_1_.

Figure 2 shows the experimental results on the dynamics of JAK2^617*F*^ mutants in the presence of TNF*α* (blue lines) and in the absence of TNF*α*. We can see that in the absence of TNF*α*, mutants retain a nearly constant fraction in the expanding population of cells. In the presence of TNF*α* the mutants are characterized by a faster growth compared to the wild type cells. Assuming that inflammation affects JAK2^617*F*^ mutants to a lesser degree compared to the wild type cells, we ask the question, what parameters are affected by inflammation? From the information on the linear growth given by figure 2, we can see that

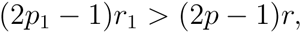

**Figure 2:**
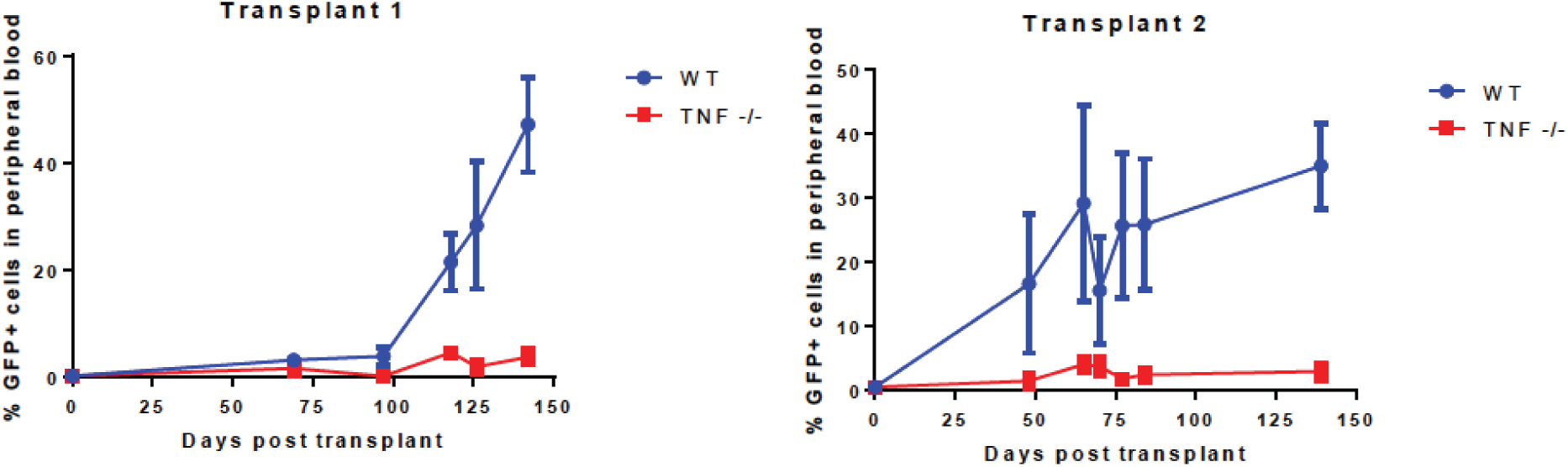
Experimentally measured growth dynamics of JAK2^617*F*^ mutants in an “empty” bone. Percentages of mutant cells with respect to the total number of cells is shown as functions of time. The blue points correspond to the stem cells transplanted in the bone of a wild type mouse; the red curves to the transplants into a TNF*α* knockout mouse. The error bars are calculated on the basis of three independent measurements. The two panels correspond to two repeats of the experiment. The initial percentage of mutants is 0.5% in all the experiments.

that is, it appears that the wild type cells experience a decrease in their division rate and/or a decrease in their proliferation probability. The latter mechanism corresponds to increased differentiation of the wild type SCs in the presence of inflammation. The mutants experience less of the effect and therefore grow faster, thus increasing in proportion.

Are these mechanisms enough to predict the invasion of the mutants under homeostatic conditions? Let us assume that in the absence of inflammation, the mutant is disadvantageous. This means that the no-inflammation case must correspond to the upper right or lower right square in the diagram in future 3. Under the influence of inflammation, only wild type parameters are affected. If in the absence of inflammation the system was in the lower right quarter of the phase space of figure 3, an increase in the probability of differentiation of wild type cells in the presence of inflammation would take the system to the left bottom quarter (extinction). Therefore, we must assume that in the absence of inflammation the system resides in the upper right quarter, and condition (34) holds. Inflammation can lead to mutant domination if the probability of wild type SC differentiation increases (that is, *P* decreases). This can happen in two possible ways: (1) Parameter *p* decreases; if *p* < 1/2 then the system transfers to the left upper quarter of the phase space (the red arrow in future 3 pointing left), otherwise a decrease in *p* must lead to the reversal of inequality (34) (the red arrow in future 3 pointing up); (2) the strength of the proliferation probability control, *q*, increases, such that inequality (34) reverses (the up arrow).

If we assume that the same mechanism is responsible for the faster growth of JAK2^617*F*^ mutants in the “empty” bone and for their selection in homeostatic, we have to conclude that it is parameter *p* that is reduced by inflammation.

### 3.5 Examples

We will use the example of the particular model in the previous section to demonstrate some general patterns derived in Section 3.3.

In particular, we will show that if inflammation influences the divisions and/or death rate of the wild type HSCs, the total number of wild type (stem and differentiated) cells will decrease under inflammation, but the percentage of HSCs may decrease or increase, depending on the parameters. To this end, we will work out two examples.

**Example 1.** The first example is obtained by using parameter values given in Table 1 in equations (28-31). As before, we consider two types of populations: wild type cells and mutant cells. Wild type cells are characterized by different sets of parameters with and without inflammation, while mutants retain the same parameters in all conditions. We can see that in this example, only parameters *p* and *r* change among the three sets. Wild type cells have division rate larger that mutants in the absence of inflammation, and smaller than mutants in the presence of inflammation. Further, the wild type cells have an increased probability to differentiate in the presence of inflammation (that is, the parameter *p* becomes smaller).

**Table 1:** Parameter values for Example 1.

| Variable | $\alpha$ | $r$ | $\omega$ | $p$ | $q$ | $D$ | $\delta$ |
| --- | --- | --- | --- | --- | --- | --- | --- |
| Wild type, no inflammation | 0.5 | 1 | 1 | 0.9 | 0.01 | 0.002 | 0.01 |
| Wild type, with inflammation | 0.5 | 0.3 | 1 | 0.6 | 0.01 | 0.002 | 0.01 |
| Mutant | 0.5 | 0.8 | 1 | 0.7 | 0.01 | 0.002 | 0.01 |

It can be shown that under these parameter values, HSCs comprise a stable equilibrium in the absence of inflammation, and they are displaced by mutants in the presence of inflammation, see figure 6 in Appendix A. Further, in the absence of mutants, the wild type cells are characterized by equilibrium values that are different depending on whether or not inflammation is present. For the parameter set in Example 1, the total population size is 6185 without inflammation, and it is 176 with inflammation, figure 8 (left) in Appendix A. Therefore, this example confirms the general trend demonstrated in section 3.3, that the total population size of wild type cells is always smaller under inflammation.

**Figure 6:**
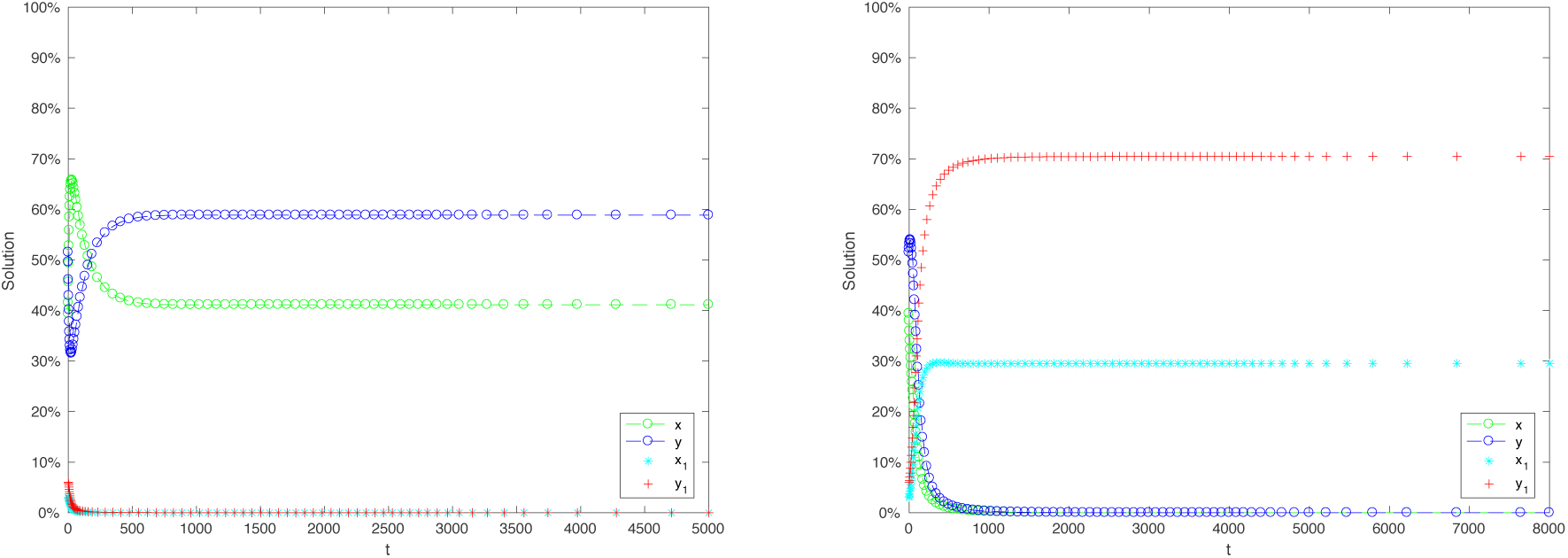
Example 1: the competition dynamics and wild type and mutant cells. Left: without inflammation, wild type cells dominate; right: with inflammation, mutant cells dominate.

**Figure 8:**
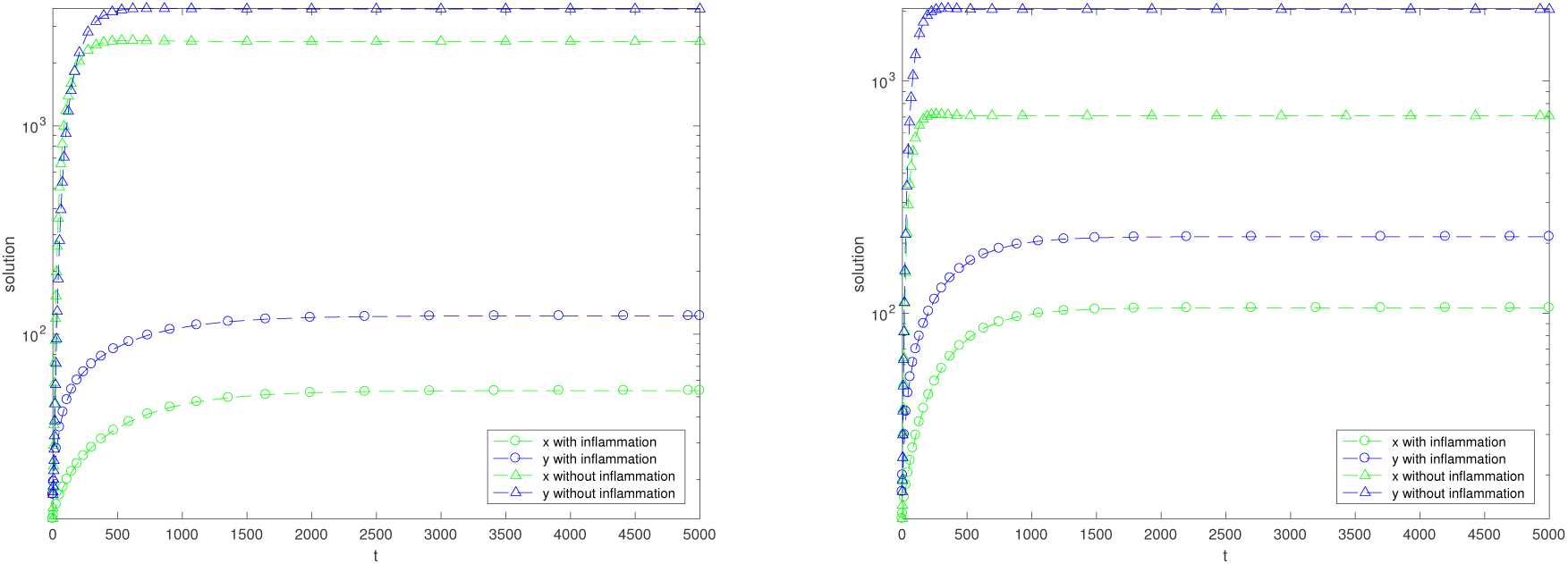
The behavior of wild type HSCs (in the absence of mutants) with and without inflammation. Left: Example 1; right: Example 2. This figure demonstrates that the total population of HSCs shrinks under inflammation.

Finally, we examine the percentage of HSCs at equilibrium with and without inflammation. Again, we assume that no mutants are present in the system (which amounts to taking *x*_1_(0) = *y*_1_(0) = 0 in the simulations. As shown in figure 4, the fraction of HSCs is 0.41 without inflammation, and it is 0.30 with inflammation. Therefore, we can see that for this particular example, the percentage of HSCs becomes smaller under inflammation.

**Figure 4:**
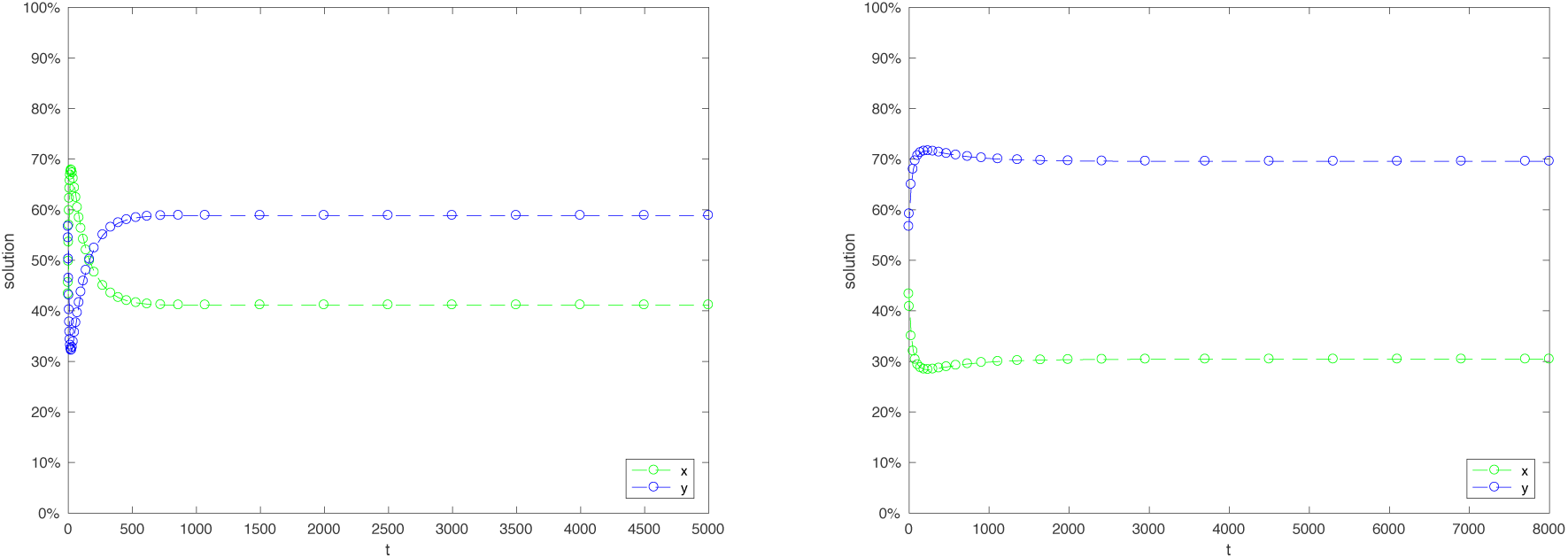
Example 1: the fraction of wild type HSCs and differentiated cells in the absence of mutants. Left: without inflammation; right: with inflammation. This figure demonstrates that the percentage of HSCs can decrease under inflammation.

**Example 2.** In the next example, we demonstrated that the percentage of HSCs can also become larger under inflammation, which is the opposite of what was observed in Experiment 1. We use parameter values specified by Table 2. As in Example 1, only two parameters are affected by inflammation, *p* and *r*, see equations (28-31).

**Table 2:** Parameter values for Example 2.

| Name | $\alpha$ | $r$ | $\omega$ | $p$ | $q$ | $D$ | $\delta$ |
| --- | --- | --- | --- | --- | --- | --- | --- |
| Wild type, no inflammation | 0.5 | 1 | 0.35 | 0.75 | 0.01 | 0.002 | 0.02 |
| Wild type, with inflammation | 0.5 | 0.26 | 0.35 | 0.6 | 0.01 | 0.002 | 0.02 |
| Mutant | 0.5 | 0.8 | 0.35 | 0.7 | 0.01 | 0.002 | 0.02 |

Again, it can be shown that HSCs comprise a stable equilibrium in the absence of inflammation, and the mutant strategy is an ESS in the presence of inflammation, figure 7 in Appendix A. When we include only wild type cells in the simulation with and without inflammation, we observe, as expected, that the steady state population is smaller under inflammation (for these parameters, it is given by 2744 and by 319 without and with inflammation respectively, figure 8 (right) in Appendix A. Again, it confirms that the total population size is smaller under inflammation. A different result is observed when we look at the percentage of HSCs in steady state, with and without inflammation. As figure 5 demonstrates, the fraction of HSCs changes from 0.26 without inflammation to 0.33 with inflammation. Therefore, for the parameters of Example 2, the cell population is enriched with HSCs under inflammation.

**Figure 5:**
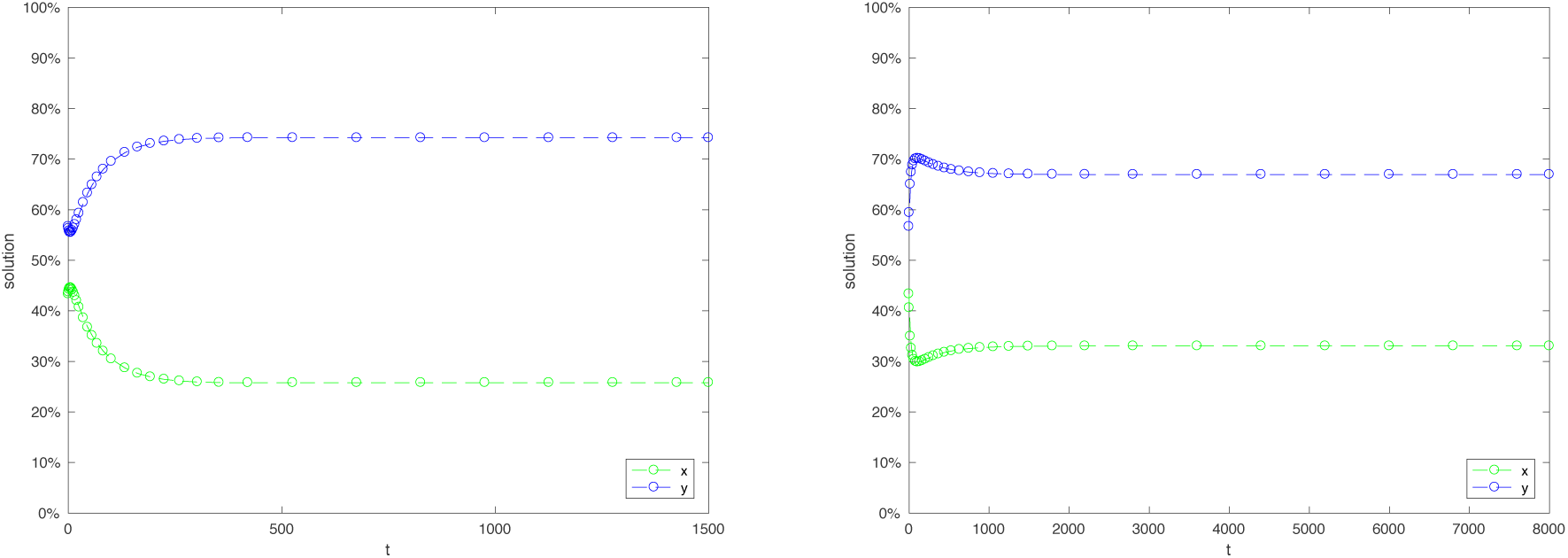
Example 2: the fraction of wild type HSCs and differentiated cells in the absence of mutants. Left: without inflammation; right: with inflammation. This figure demonstrates that the percentage of HSCs can increase under inflammation.

**Figure 7:**
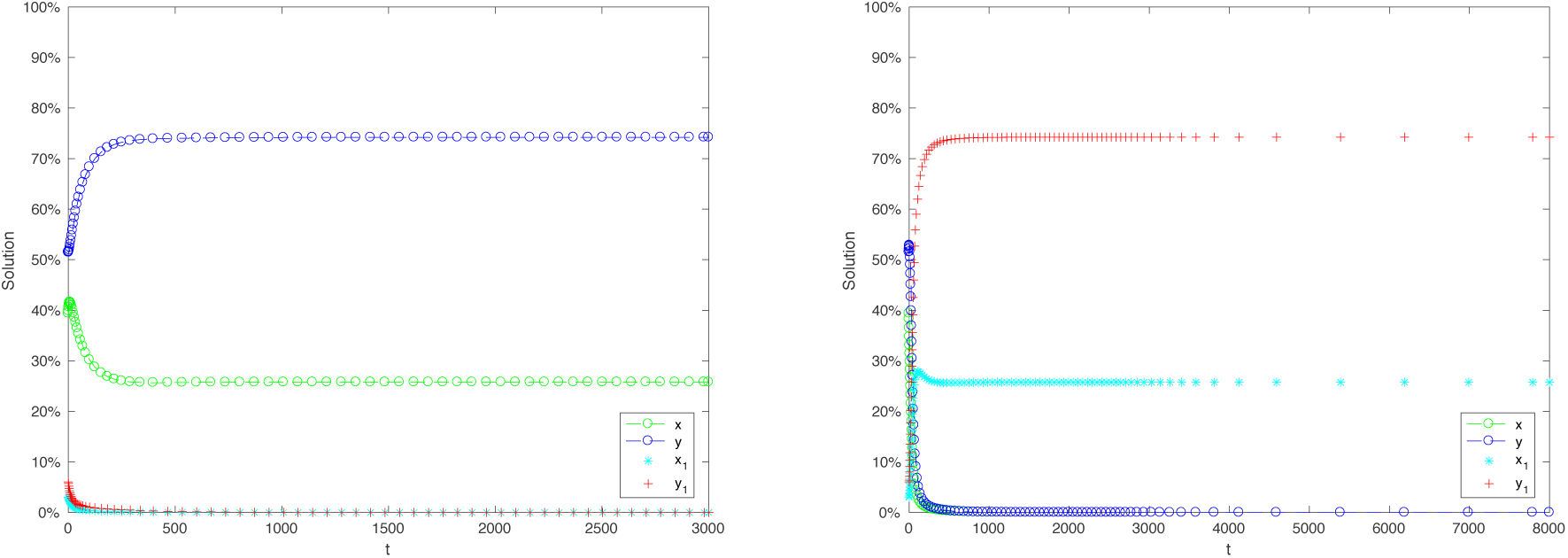
Example 2: the competition dynamics and wild type and mutant cells. Left: without inflammation, wild type cells dominate; right: with inflammation, mutant cells dominate.

## 4 Discussion

We have created a model of HSC dynamics in the context of inflammation and its role in MPN, mediated by JAK2^617*F*^ mutants. Because of significant uncertainties that exist in the mechanisms governing the different cellular kinetic rates, we chose to implement an axiomatic approach. This means that we spell out the exact mathematical assumptions that follow from our understanding of the biological system, and investigate what predictions we can make with the resulting model, without implementing arbitrary (and unknown) functional forms. We have looked specifically at the competition dynamics of wild type and mutant HSCs in the absence and in the presence of inflammation. It turned out that, surprisingly, even a very general model gave rise to a number of stable predictions. They are listed as follows:

1. In the absence of HSC death, the evolutionary advantage of mutant cells can only be conferred by an increase in differentiation probability of HSCs in the presence of inflammation
2. If the differentiation probability of HSCs in the presence of inflammation is increased, the numbers of HSC and differentiated cells decreases at equilibrium (in the absence of mutants), and the percentage of HSCs also decreases.
3. In the presence of HSC death, the evolutionary advantage of mutant cells can be conferred either by an increase in differentiation probability of HSCs, or by an increase of their division-to-death ratio, in the presence of inflammation.
4. In this case, while the number of cells is decreased at the equilibrium, the percentage of HSCs may decrease or increase, depending on the particular properties of the control network governing HSC fate decisions.

These predictions have been confirmed with particular numerical models, which are specific implementations of the general modeling approach.

The result obtained here may be useful not only for studying MPN but also for other biomedical scenarios. MPN is an ideal system to investigate the role of inflammation in aberrant cell dynamics, for the following reasons: (1) MPN is a relatively genetically homogenous disease, with JAK2^617*F*^ as the driver mutation in the majority of patient, this makes it an ideal disease to study the dynamics of clonal evolution in hematologic malignancies. (2) Mouse models for MPN exist that recapitulate the human phenotype and can be easily studied. Many other cancers, for example colon cancer, do not have an adequate mouse model. (3) There is a strong indication that inflammation not only plays a role in MPN, but inflammation likely exerts several types of influence in this disease, internal and external, both of which will be studied. (6) Findings in MPN could then be applied to other hematologic malignancies and potentially solid tumors.

The value of using MPN as our model system goes well beyond its utility as a good representation of clonal evolution in hematologic malignancies. There is a real need for interventions that actually impact disease progression in MPN. There are currently no therapies other than bone marrow transplantation that impact disease progression in MPN, and bone marrow transplantation is a reasonable option for only a minority of MPN patients. The current therapeutic goals in MPN are simply to reduce the risk of blood clots and manage symptoms. Interventional clinical trials in MPN focus on end stage disease, with minimal attention to patients with early stage disease. However, we believe that intervening in early stage MPN will make the most meaningful impact on patient outcomes. An understanding of the conditions that promote the emergence of MPN as well as progression from early stage to late stage disease is vital for designing chemoprevention measures that could halt progression of disease in patients with early stage MPN or could prevent disease in healthy individuals at risk of developing MPN. Recent research strongly indicates that inflammatory processes create an environment that promotes the selection of JAK2^617*F*^ mutant MPN cells and that interference with these inflammatory processes can prevent the expansion of these mutant clones (Fleischman et al, 2011). Details of how this selection occurs, however, are not well understood. The current paper creates a theoretical framework for studying these issues.

This paper identifies several patterns compatible with currently available measurements. The next step is to create more detailed models, based on experimental research, to be able to make further, mode detailed, testable predictions.

## Acknowledgements

DW and NK gratefully acknowledge the support of NIH grant 1 U01 CA187956-01. AF is supported by V Foundation Scholar Award and DoD Career Development Award CA150493.

## A Additional figures detailing the dynamics of Examples 1 and 2

Here we provide additional information about the simulations performed for Examples 1 and 2 in the main text. Figures 6 and 6 demonstrate the basic settings. In both examples, the wild type cells dominate in the absence of inflammations (left panels) and they are displaced by mutant cells in the presence of inflammation (right panels) by competitive exclusion. In all these simulations, we include nonzero values for all four cell types as initial conditions. In figure 8 we study the dynamics of wild type cells alone, in the absence and in the presence of inflammation. The left (right) panel of this figure corresponds to parameters of Example 1 (Example 2). In the simulations for this figure, we start with zero initial values of *x*_1_ and *y*_1_, that is, no mutants are included.

